# Destruction of DNA-binding proteins by programmable O’PROTAC: Oligonucleotide-based PROTAC

**DOI:** 10.1101/2021.03.08.434493

**Authors:** Jingwei Shao, Yuqian Yan, Donglin Ding, Dejie Wang, Yundong He, Yunqian Pan, Wei Yan, Anupreet Kharbanda, Hong-yu Li, Haojie Huang

**Affiliations:** Department of Pharmaceutical Sciences, College of Pharmacy, University of Arkansas for Medical Sciences, Little Rock, AR 72205, USA; Department of Biochemistry and Molecular Biology, Mayo Clinic College of Medicine and Science, Rochester, MN 55905, USA; Department of Urology, Mayo Clinic College of Medicine and Science, Rochester, MN 55905, USA; Mayo Clinic Cancer Center, Mayo Clinic College of Medicine and Science, Rochester, MN 55905, USA

**Author notes:** These authors contributed equally (Jingwei Shao, Yuqian Yan, Donglin Ding). **Corresponding authors**, Hong-yu Li and Haojie Huang.

## Abstract

DNA-binding proteins including transcription factors (TFs) play essential roles in gene transcription and DNA replication and repair during normal organ development and pathogenesis of diseases such as cancer, cardiovascular disease and obesity, deeming to be a large repertoire of attractive therapeutic targets. However, this group of proteins are generally considered undruggable as they lack an enzymatic catalytic site or a ligand binding pocket. PROteolysis-TArgeting Chimera (PROTAC) technology has been developed by engineering a bifunctional small molecule chimera to bring a protein of interest (POI) to the proximity of an E3 ubiquitin ligase, thus inducing the ubiquitination of POI and further degradation through proteasome pathway. Here we report the development of Oligonucleotide-based PROTAC (O’PROTACs), a class of noncanonical PROTACs in which a TF-recognizing double-stranded oligonucleotide is incorporated as a binding moiety of POI. We demonstrate that O’PROTACs of ERG and LEF1, two highly cancer-related transcription factors selectively promote degradation of these proteins and inhibit their transcriptional activity in cancer cells. The programmable nature of O’PROTACs indicates that this approach is applicable to destruct other TFs. O’PROTACs not only can serve as a research tool, but also can be harnessed as a therapeutic arsenal to target DNA binding proteins for effective treatment of diseases such as cancer.

## Introduction

A large group of DNA-binding proteins act as transcription factors (TFs) that transcriptionally activate or suppress gene expression by interacting with specific DNA sequence and transcription co-regulators. Approximately 2,000 TFs have been identified in eukaryotic cells and they are associated with numerous biological processes. Among them, approximately 294 TFs are associated with cancer development, which account for ~19% of oncogenes^1^. Therefore, targeting TFs associated with cancer development appears to be an appealing strategy for cancer treatment.

In the last decades, small molecule modulators have been developed to target nuclear receptors on the basis that this class of TFs contains a clearly defined ligandbinding pocket^2^. However, most of other TFs are difficult to target because they lack a ligand binding pocket. As the knowledge regarding the mechanisms of the assembly of transcription complexes has increased exponentially, different strategies to modulate the activity of TFs with small molecule compounds have emerged, including blocking protein/protein interactions, protein/DNA interactions, or chromatin remodeling/epigenetic reader proteins^3^. However, the development of traditional small molecules inhibiting non-ligand TFs remains very challenging, and a new targeting strategy to overcome the hurdle is highly demanded.

PROTACs are heterobifunctional small molecules composed of a POI ligand as a warhead, a linker and an E3 ligase ligand. The PROTAC molecule recruits the E3 ligase to the POI and induced the ubiquitination of the latter and further degradation by the proteasome pathway. PROTAC technology has greatly advanced during the last decade. It has been proved that PROTACs are capable of degrading a variety of proteins, including enzymes and receptors^4–7^. Two PROTACs, ARV-110 and ARV-471 which are androgen receptor (AR) and estrogen receptor (ER) degraders, respectively have entered phase I clinical trials^8–9^. PROTACs offer several advantages over small molecule inhibitors including expanding target scope, improving selectivity, reducing toxicity and evading inhibitor resistance^10^. This suggests that PROTAC technology is a new promising modality to tackle diseases, in particular for cancer. Most recently, PROTACs have been designed to degrade TFs. Wang’s group developed a potent and signal transducers and activators of transcription 3 (STAT3)-specific degrader based on an STAT3 inhibitor SI-109 and demonstrated its targeting efficacy *in vivo*^11^. Crews’ group reported the development of Transcription Factor Targeting Chimeras (TRAFTACs)^12^, which utilize haloPROTAC, dCas9-HT7 and dsDNA/CRISPR-RNA chimeras to degrade TFs. Nevertheless, this approach uses the artificially engineered dCas9-HT7 fusion protein as a mediator, which limits its potential use in clinic.

ERG transcription factor belongs to the ETS family and is involved in bone development, hematopoiesis, angiogenesis, vasculogenesis, inflammation, migration and invasion^13^. Importantly, it is overexpressed in approximately 50% of all human prostate cancer cases including both primary and metastatic prostate cancer due to the fusion of *ERG* gene with the androgen-responsive *TMPRSS2* gene promoter^14^. *TMPRSS2-ERG* gene fusion results in aberrant overexpression of truncated ERG, implying that increased expression of ERG is a key factor to drive prostate cancer progression^15^. Therefore, therapeutic targeting ERG is urgently needed to effectively treat prostate cancer patients. Lymphoid enhancer-binding factor 1 (LEF1) is another highly cancer-related TF. It belongs to T cell factor (TCF)/ LEF1 family. Complexed with β-catenin, LEF1 promotes the transcription of Wnt target genes^16^. LEF1 also can facilitate epithelial-mesenchymal transition (EMT)^17^. Aberrant expression of LEF1 is implicated in several cancer types and related to cancer cell proliferation, migration, and invasion^18^. Hence, LEF1 is another ideal target for cancer treatment.

In the present study, we introduce a new strategy to target TFs using O’PROTACs, in which a double-stranded oligonucleotide is incorporated as POI binding moiety in PROTAC (Figure 1). We demonstrate that ERG O’PROTAC promotes proteasomal degradation of ERG protein and inhibits ERG transcriptional activity. Akin to ERG degrader, LEF1 O’PROTAC induces the degradation of LEF1. Consequently, its target gene expression and prostate cancer cell growth was also effectively inhibited.

**Fig 1.**
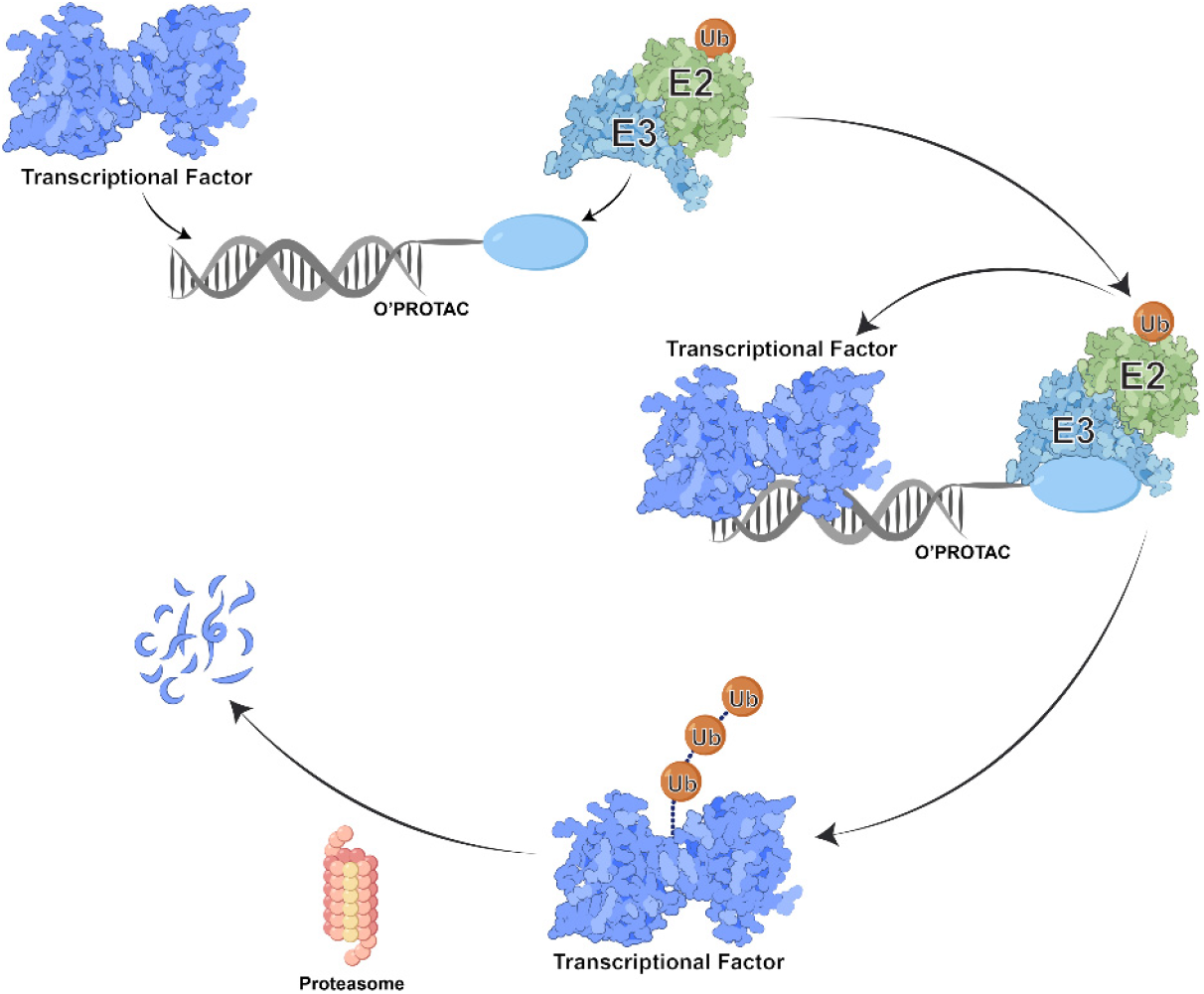
Schematic depicting the working principle of O’PROTAC.

## Results

### Design of O’PROTACs

ERG recognizes a highly conserved DNA binding consensus sequence including the 5’-GGAA/T-3’ core motif^19^. We designed a 19-mer double-stranded oligonucleotide containing the sequence of ACGG<u>ACCGGAAAT</u>CCGGTT with the ERG binding moiety underscored. As for the E3 ligase-recruiting element, we selected the widely used pomalidomide and VHL-032, which are capable of hijacking cereblon and von Hippel-Lindau (VHL) respectively. PROTAC exerts its function based on the formation of ternary complex, in which a linker plays an important role.

Therefore, we designed and synthesized six phosphoramidites with different linkers in different lengths and types, three of which are linked to pomalidomide and three with VHL-032 (**P1-6**, **Table 1**). The phosphoramidite was attached to the 5’ terminal of one DNA strand through DNA synthesizer. After annealing, we generated six O’PROTACs (OPs) for both ERG and LEF1. (**Table S1**).

**Table 1.**
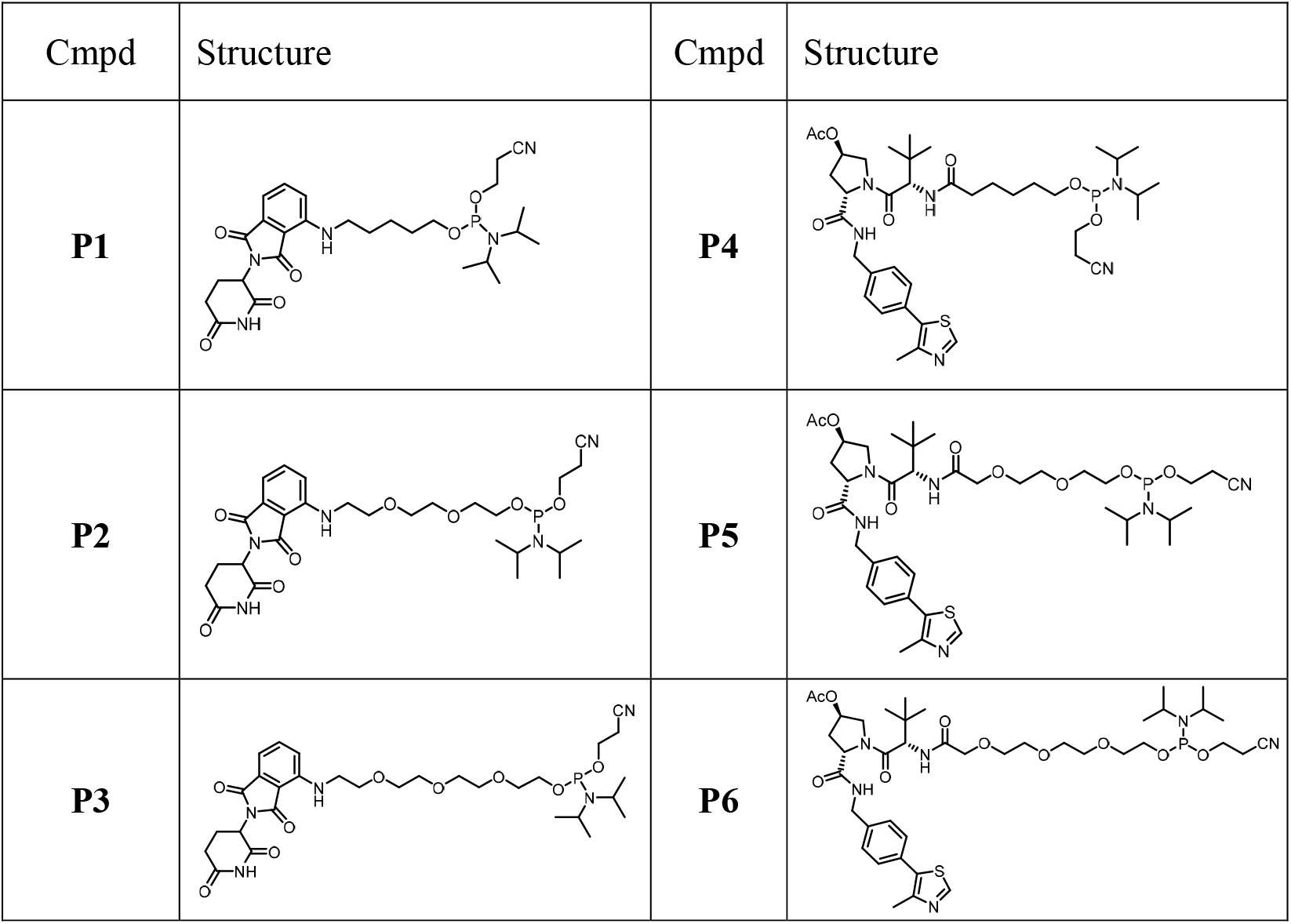
The structures of phosphoramidites P1-6.

### Chemical synthesis of P1-6

The synthesis of **P1-6** was illustrated in Scheme 1. 4-Fluoro-thalidomide and VHL-032 were prepared according to literature procedures^20–21^. The straightforward nucleophilic aromatic substitution reaction of 4-fluoro-thalidomide with different amines provided key intermediates **8a-c**. VHL-032 was coupled with various carboxylic acids containing TBDPS protected hydroxyl group to deliver intermediates **8d-f.** Subsequent acetylation of the hydroxyl groups in **8d-f** and removal of the TBDPS protection produced intermediates **10a-c**. Phosphitylation of **8a-c** or **10a-c** with Cl-POCEN^*i*^Pr_2_ yielded **P1-6** in the presence of DIPEA.

**Scheme 1.**
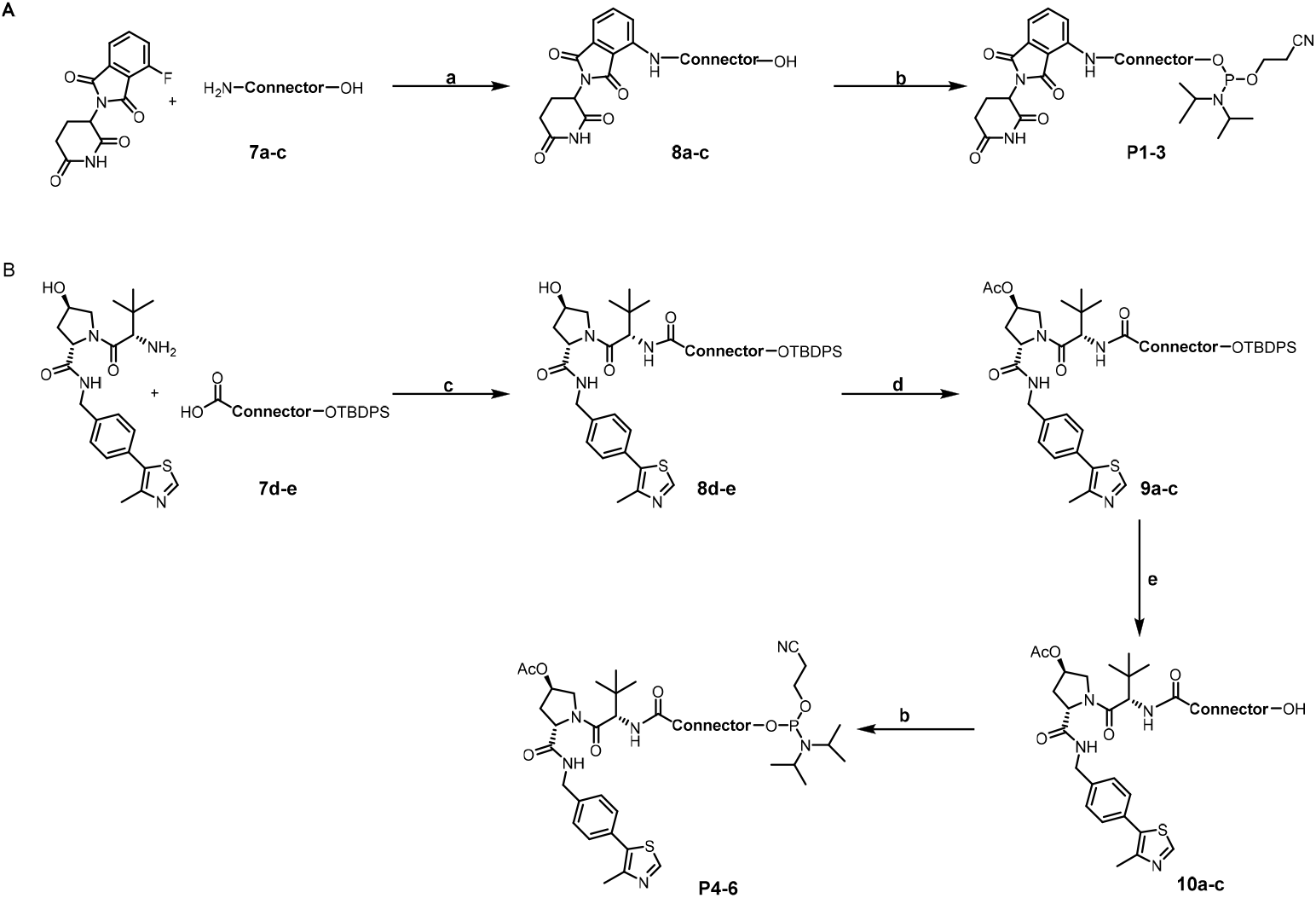
Synthesis of P1-6*a*. ^*a*^Reagents and conditions: (a) DIPEA, NMP, MW, 100 °C, 3 h; (b) Cl-POCEN^*i*^Pr_2_, DIPEA, DCM, 1 h, rt. (c) HATU, TEA, DMF; (d) Ac2O, DMAP, DCM, 1 h; (e) TBAF, THF.

### ERG O’PROTACs promote proteasome degradation of WT and TMPRSS2-ERG proteins

The nucleic acid-based agents typically rely on lipid-mediated transfection to deliver them into cells. FITC-labelled ERG O’PROTAC was synthesized to determine the transfection efficiency under a fluorescent microscope. We transfected 293T cells with 100 or 1,000 nM of O’PROTAC with or without lipofectamine 2000. As expected, the presence of lipofectamine greatly enhanced the cellular uptake comparing with mock transfection (Figure 2A). However, there was no difference in uptake efficacy between low (100 nM) and high concentration (1,000 nM) (Figure 2A), probably owing to the saturation of the positively charged lipid with negatively charged oligonucleotide.

**Figure 2.**
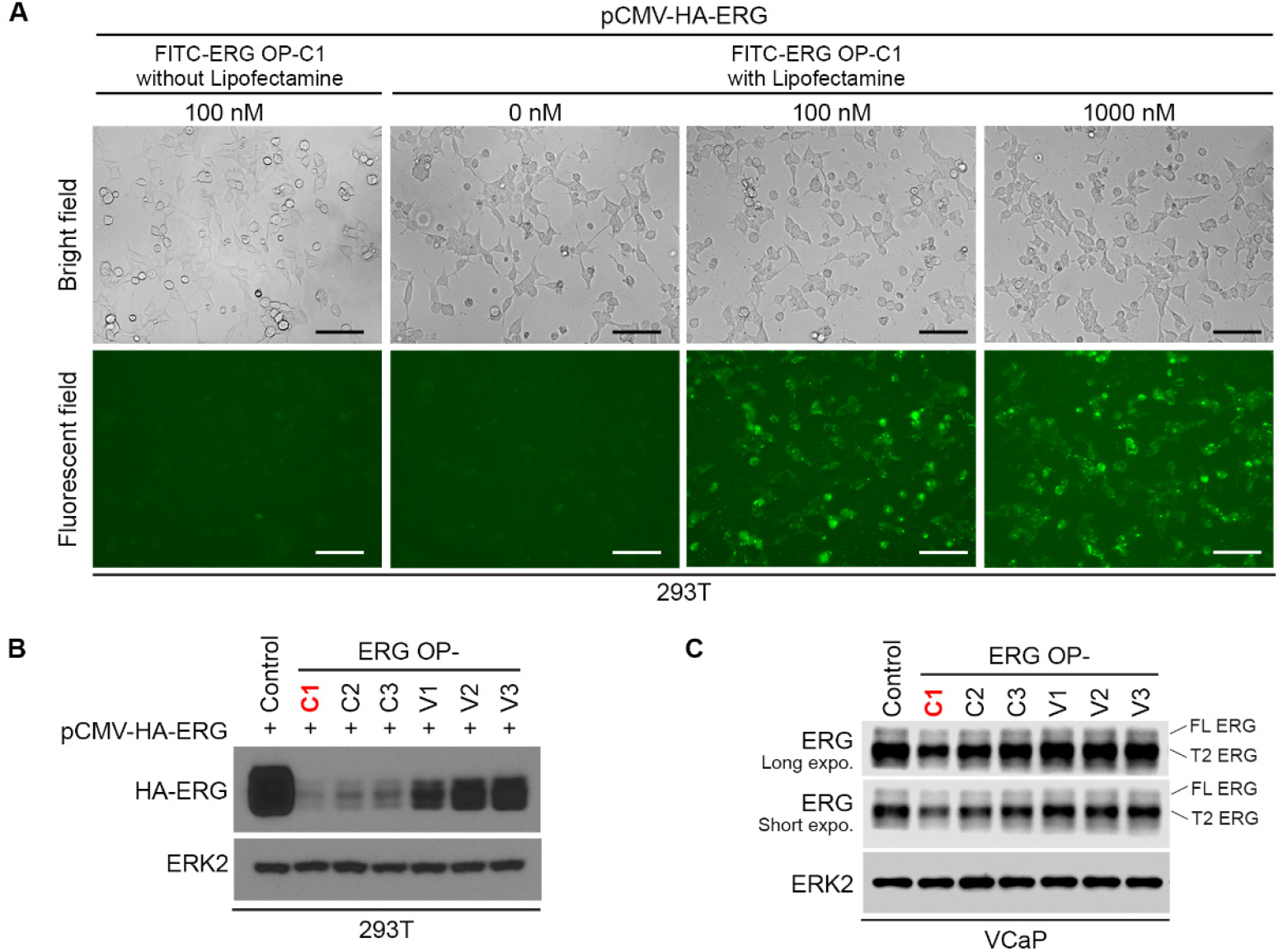
ERG O’PROTACs degrade ERG protein in cultured cells. (A) 293T cells were transfected with FITC-labelled ERG O’PROTAC-13 (100 nM and 1,000 nM) and the transfection efficiency was monitored 48 h post-transfection using a fluorescent microscope. Scale bar: 50 μm. (B) 293T cells were transfected with HA-ERG plasmid and a control or six indicated ERG O’PROTACs (100 nM) and harvested for western blot analysis 48 h post-transfection. ERK2 was used as a loading control. (C) VCaP cells were transfected with a control or six indicated ERG O’PROTACs (100 nM) and cells were harvested for western blot analysis 48 h posttransfection. Both endogenous full-length (FL) wild-type and TMPRSS2-ERG (T2-ERG) truncated ERG were detected.

To assess the effects of ERG O’PROTACs on ERG proteins in cells, 293T cells were transfected with exogenously expressing HA-ERG plasmid and six ERG O’PROTACs at 100 nM for 48 hours and ERG protein level was measured by western blot. A significant decrease in ERG protein level was observed upon treatment with ERG OP-C1-3 attached with pomalidomide while the effects of ERG OP-V1-3 conjugated with VHL-032 were much modest (Figure 2B). To further demonstrate the cellular effect on endogenous ERG protein level, we tested ERG O’PROTACs in ERG-overexpressed human prostate cancer cell line VCaP which expresses both full-length ERG and TMPRSS2-ERG truncation^22^. Similar to the effect on ectopically expressed ERG, ERG OP-C1-3 also effectively decreased endogenous ERG protein in VCaP cells (Figure 2C). These data also imply that a shorter linker such as five carbon atoms (ERG OP-C1) might favorably form a more stable ternary complex. Although ERG OP-C1 significantly decreased ERG protein level, proteinase inhibitor MG132 blocked this degradation (Figure 3), suggesting ERG O’PROTAC degrades ERG protein via proteasome pathway.

**Figure 3.**
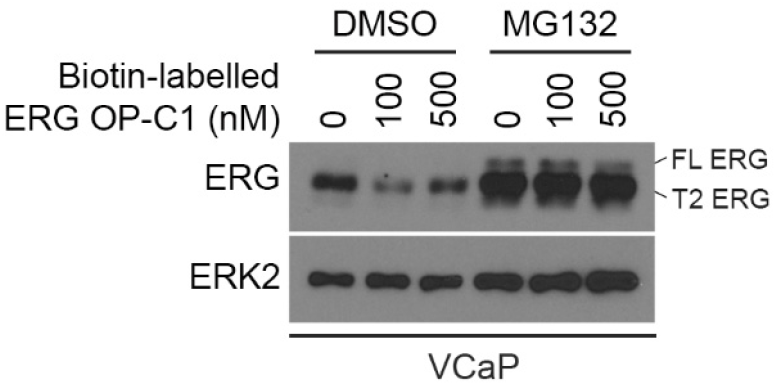
ERG O’PROTAC promotes ERG degradation via the protostome pathway. VCaP cells were transfected with increasing concentrations of ERG **OP-C1** for 36 h, followed by treatment of the proteinase inhibitor MG132 for 12 h and western blot analysis.

In vitro biotin pulldown assay showed that a significant amount of HA-ERG expressed in 293T cells was pulled down by biotin-labelled ERG OP-C1 and OP-C2 (Figure 4), indicating that these two O’PROTACs strongly interact with ERG protein. This result also provides a plausible explanation for the better effect of these two O’PROTACS on ERG degradation.

**Figure 4.**
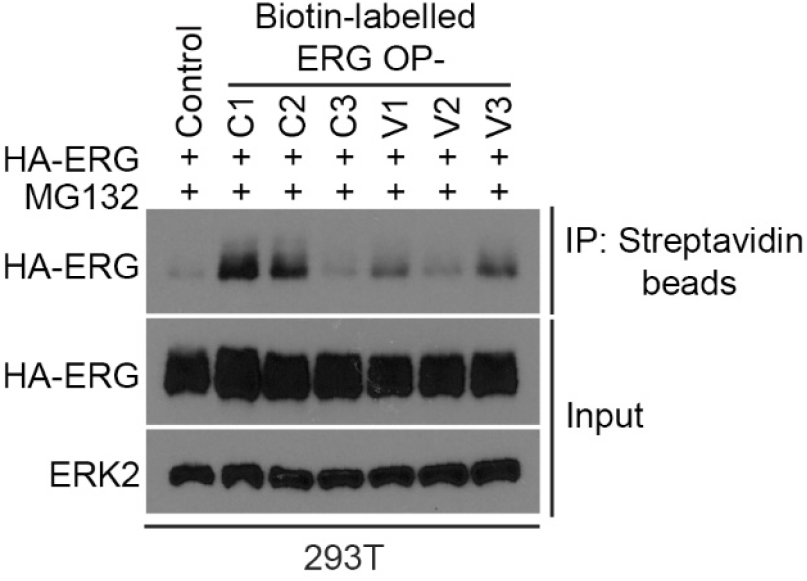
ERG O’PROTACs bind to ERG. 293T cells were transfected with HA-ERG plasmid in combination with control (non-biotin labelled) or six indicated biotin-labelled ERG O’PROTACs (100 nM) and harvested for anti-biotin (streptavidin) pull-down assay 48 h post-transfection.

Time-course studies showed that ERG O’PROTACs took effects starting from 12 hours until 48 hours examined (Figure 5A). Consistent with the finding in 293T cells (Figure 2A), the dose-course experiments revealed that 100 nM of ERG OP-C1 showed a significant inhibition of ERG protein level and this effect was not improved by higher concentrations such as 500 and 1,000 nM, indicating that ERG OP-C1 is probably saturated in a higher concentration (Figure 5B). Additionally, treatment of VCaP cells with ERG OP-C1 inhibited mRNA expression of ERG target genes including *ADAM19, MMP3, MMP9, PLAT* and *PLAU* (Figure 5C), suggesting that ERG O’PROTAC inhibits ERG transcriptional activity in VCaP prostate cancer cells.

**Figure 5.**
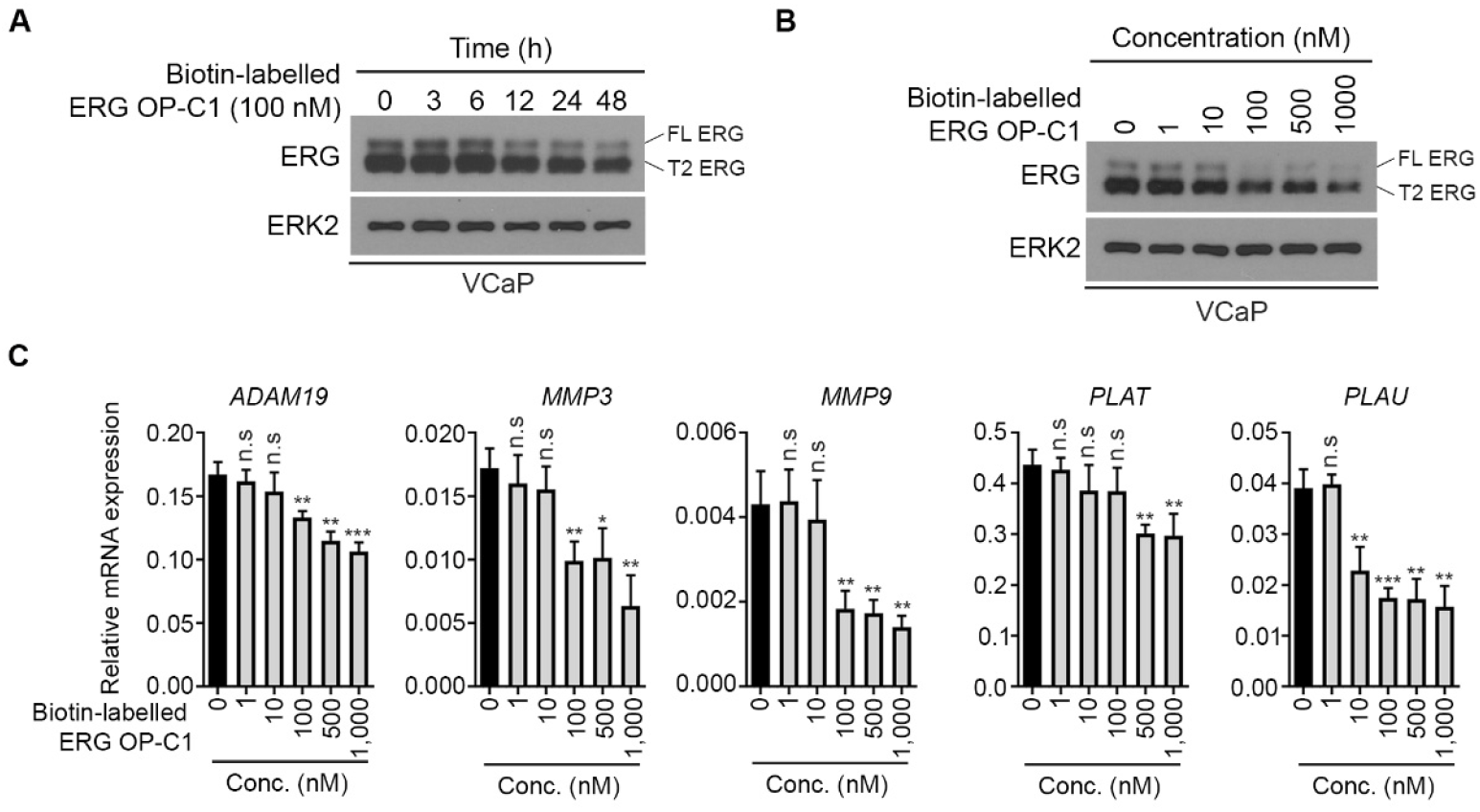
ERG O’PROTAC inhibits ERG transcriptional activity. (A) VCaP cells were transfected with 100 nM of biotin-labelled ERG **OP-C1**. Cells were harvested at the different time points followed by western blot analysis. (B and C) VCaP cells were transfected with different concentrations of biotin-labelled ERG **OP-C1** and harvested 45 h post-transfection for western blot analysis (B) and RT-qPCR analysis of mRNA expression of the indicated ERG-targeted genes *(ADAM19, MMP3, MMP9, PLAT* and *PLAU). P* values were calculated using the unpaired two-tailed Student’s *t*-test; * *P* < 0.05; ** *P* < 0.01; *** *P* < 0.001, n.s., not significant.

### Targeting other TFs for degradation by O’PROTACs

To extend the utility of O’PROTACs, we turned to another transcription factor LEF1. LEF1 acts as a DNA binding subunit in the β-catenin/LEF1 complex and exerts transcriptional regulation via binding to the nucleotide sequence 5’-A/TA/TCAAAG-3’ ^23^. We designed 18-mer double-stranded oligonucleotide containing the sequence of TACAAAGATCAAAGGGTT as the LEF1 binding moiety. Six LEF1 O’PROTACs (**Table S1**) were synthesized using the same protocol as for the ERG O’PROTACs.

We first evaluated the degradation capability of each LEF1 O’PROTACs in PC-3 prostate cancer cell line. Western blot assay was utilized to detect the expression of LEF1 protein. As shown in Figure 6, LEF1 OP-V1 potently induced LEF1 degradation in PC-3 cells at a lower concentration (100 nM) while other LEF1 O’PROTACs were less or not active. This result is similar with ERG O’PROTACs, suggesting that both linker length and E3 ligase are important factors for degradation of a specific TF.

**Figure 6.**
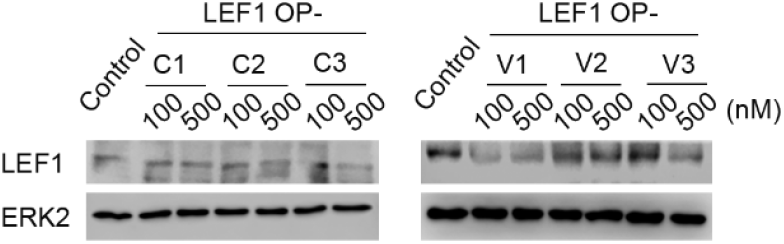
LEF-1 O’PROTACs degrade LEF1 protein in cultured cells. PC-3 cells were transfected with a control (500 nM) or six indicated LEF1 O’PROTAC at different concentrations (100 and 500 nM) and cells were harvested for western blot analysis 48h post-transfection. ERK2 was used as a loading control.

Next, we examined the effect of LEF1 O’PROTAC on the transcriptional activity of the β-Catenin/LEF1 complex. We found that treatment of PC-3 prostate cancer cells with LEF1 OP-V1 downregulated mRNA expression of *CCND1* and *c-MYC*, two target genes of β-Catenin/LEF1 in a dose-dependent manner (Figure 7A and B). While LEF1 OP-V1 treatment did not affect mRNA expression of *LEF1* gene, it markedly decreased expression of LEF1 and its target protein Cyclin D1 at the protein level in PC-3 (Figure 7A). Importantly, LEF1 OP-V1 significantly inhibited PC-3 cell growth in a time- and dose-dependent fashion (Figure 7A and C). Similar results were obtained in another prostate cancer cell line DU145 (Figure 7D-F). Collectively, LEF1 OP-V1 is a potent LEF1 degrader.

**Figure 7.**
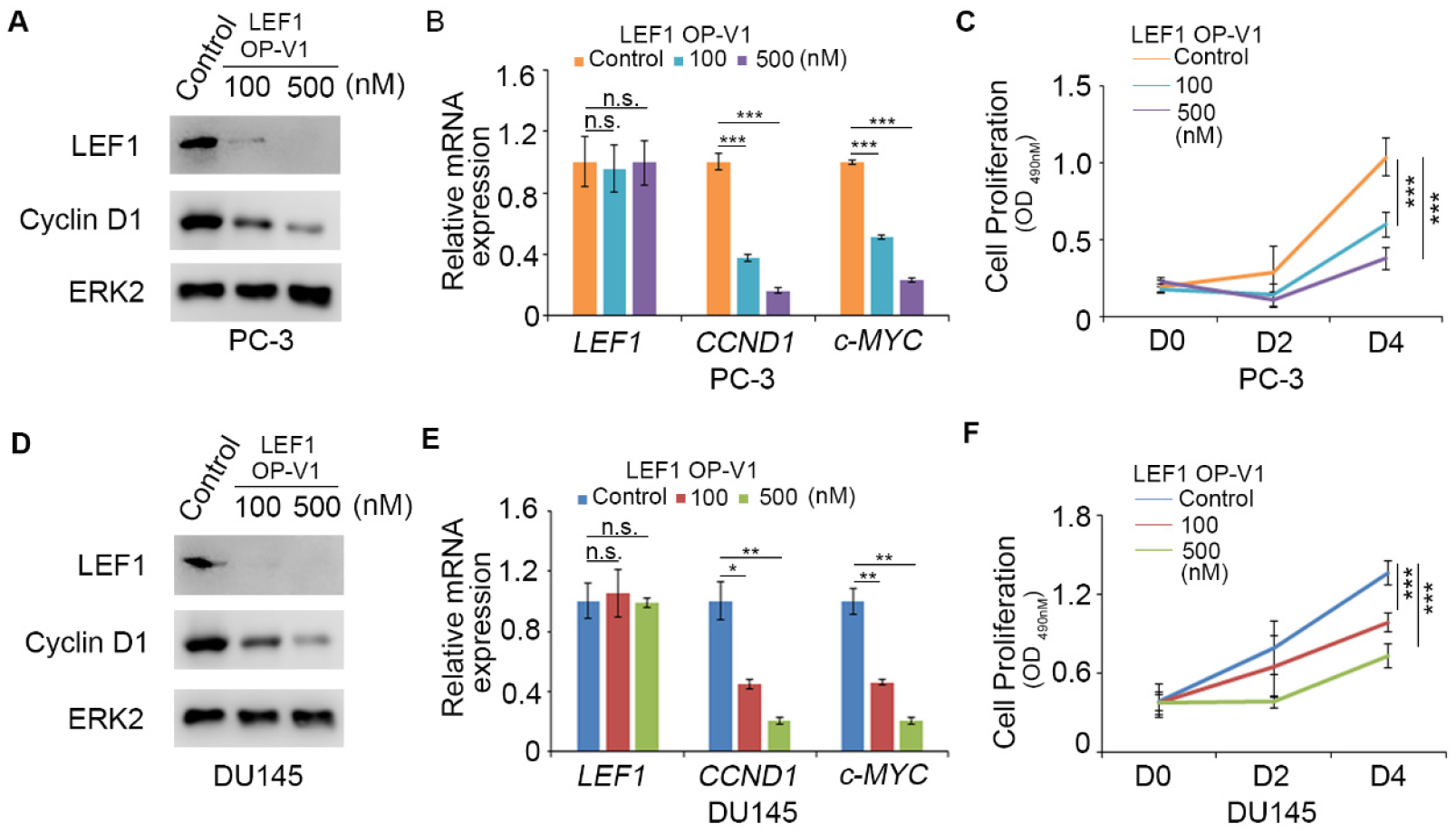
LEF1 O’PROTAC inhibits LEF1 target gene expression and prostate cancer cell proliferation. (A-C) PC-3 cells were transfected with a control (500 nM) or different concentrations of LEF1 OP-V1. At 48 h post-transfection cells were harvested for western blot analysis (A), RT-qPCR analysis of mRNA expression of LEF1 targeted genes (*CCND1* and *c-MYC*) (B), and MTS assay at different days after treatment (C). (D-F) DU145 prostate cancer cells were transfected with a control (500 nM) or different concentrations of LEF1 OP-V1. Transfected cells were subjected to western blot (D), RT-qPCR (E) and MTS assay (F). *P* values were calculated using the unpaired two-tailed Student’s *t*-test; * *P* < 0.05; ** *P* < 0.01; *** *P* < 0.001, n.s., not significant.

## Discussion

In this study we take a new strategy of degrading “undruggable” transcription factors by employing O’PROTACs. O’PROTAC exploits natural “ligand” of transcription factors, namely specific DNA sequence, attached to an E3 ligase ligand via a linker. The tactic has been successfully applied to degrade ERG and LEF1 TFs with potent efficacy in cultured cells.

Conventional PROTAC technology is rapidly evolving with some of them are in clinical trials; however, it inherits certain limitations. First, most of the reported PROTACs rely on the existing small molecules as POI targeting warhead, which make it difficult to be applied to “undruggable” targets like TFs. Additionally, due to their high molecular weight (600~1400 Da), PROTACs suffer from poor cell permeability, stability and solubility^24^. In comparison with classic small molecule drugs, PROTACs are significantly less druggable. O’PROTACs hold enormous potentials to transcend the limitations of conventional PROTACs. Because of their modalities, degraders can be rationally programmed according to the DNA binding sequence of a given TF, thus theoretically making it possible to target any TF of interest. Our data suggest that the efficacy of O’PROTACs can be further optimized by altering the lengths and types of a linker and the E3 ligase ligand. Moreover, the synthesis of O’PROTAC is highly simple and efficient, which facilitates the rapid development of a O’PROTAC library for high-throughput screening of the most potent TF degraders. O’PROTAC could be applied to any proteins bound to DNA duplexes.

Hall and colleagues recently report RNA-PROTACs, which utilize singlestranded RNA (ssRNA) to recruit RNA-binding protein (RBP). The binding of RBP with RNA heavily relies on both sequence motif and secondary structure^25–26^. Predicting the interaction between RNA and RBP is challenging due to the high flexibility of RNA^27–28^. However, double-stranded DNA bear a well-defined three-dimensional duplex structure; therefore, the protein binding region is accessible and predictable. Hence, O’PROTAC is programmable by changing the nucleotide sequence that binds protein. Additionally, compared with double-stranded oligonucleotide, ssRNA is susceptible to deleterious chemical or enzymatic attacks^28^. Taken together, O’PROTAC is desirable due to its readily predictability and superior stability.

Oligonucleotide drug development has become a main stream for new drug hunting in the last decade^29^. The catalytic advantage of PROTACs^30^ incorporated into oligonucleotide drugs could further fuel the field. Moreover, the delivery of oligonucleotide drugs has been advanced significantly in the recent years, notably for mRNA COVID-19 vaccine^31–32^. Therefore, O’PROTACs can be a complementary drug discovery and development platform to conventional PROTACs to derive clinical candidates and accelerate drug discovery.

