## Supplementry Information for "Destruction of DNA-binding proteins by programmable O’PROTAC: Oligonucleotide-based PROTAC"

*** Corresponding authors**

### EXPERIMENTAL SECTION

#### Synthesis of phosphoramidites 1-6

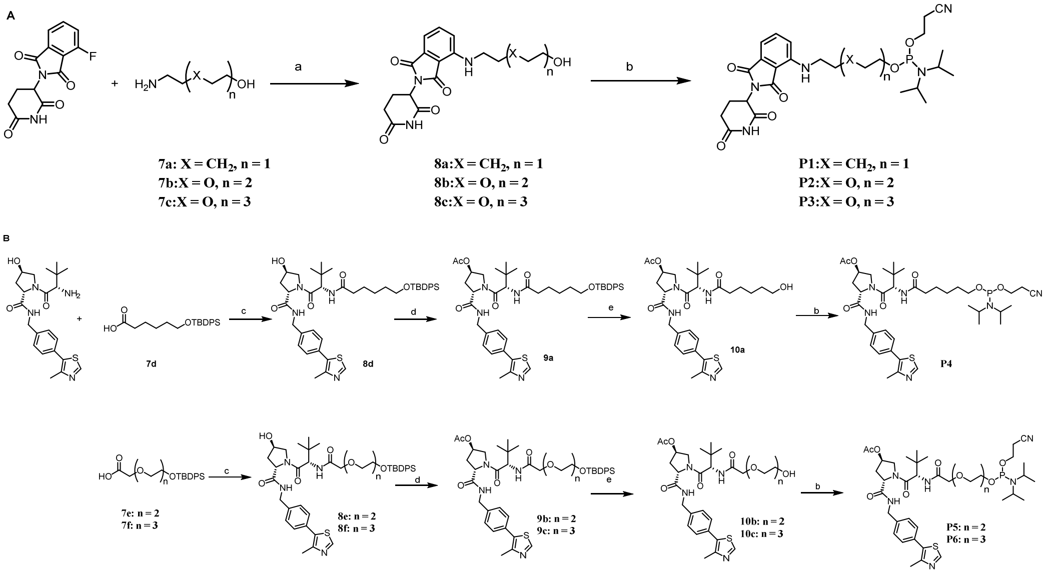


[a]Reagents and conditions: (a) DIPEA, NMP, MW, 100 °C, 3 h; (b) Cl-POCEN$^i$Pr$_2$, DIPEA, DCM, 2 h, rt. (c) HATU, TEA, DMF, rt; (d) $Ac_2O$, DMAP, DCM, 1 h; (e) TBAF, THF, rt.

**Synthesis of compound 8a-c:** Compound 4-fluoro-thalidomide (1.0 equiv) was dissolved in NMP, DIPEA (2.0 equiv) and **7a-c** (1.5 equiv) were added, the mixture was heated to 100 °C under microwave condition for 3 hours. then the mixture was absorbed on diatomite and purified by reversed-phase flash chromatography ($H_2O$: MeOH=90:10 to 50:50), giving compounds **8a-c**.

**2-(2,6-dioxopiperidin-3-yl)-4-((5-hydroxypentyl)amino)isoindoline-1,3-dione (8a):** Yellow solid, 65%.$^1$H NMR (400 MHz, $CDCl_3$) δ 8.16 (s, 1H), 7.49 (dd, $J$ = 8.5, 7.1 Hz, 1H), 7.09 (d, $J$ = 7.1 Hz, 1H), 6.88 (d, $J$ = 8.5 Hz, 1H), 4.91 (dd, $J$ = 12.1, 5.4 Hz, 1H), 3.66 (q, $J$ = 6.3 Hz, 2H), 3.28 (t, $J$ = 7.0 Hz, 2H), 2.93 – 2.67 (m, 3H), 2.12 (ddd, $J$ = 9.6, 5.8, 2.9 Hz, 1H), 1.75 – 1.66 (m, 2H), 1.64 – 1.59 (m, 2H), 1.54 – 1.46 (m, 2H).

**2-(2,6-dioxopiperidin-3-yl)-4-((2-(2-(2-hydroxyethoxy)ethoxy)ethyl)amino)isoindoline-1,3-dione (8b)**: Yellow oil, 40%. $^1$H NMR (400 MHz, $CDCl_3$) δ 8.31 (s, 1H), 7.48 (dd, *J* = 8.5, 7.2 Hz, 1H), 7.10 (d, *J* = 7.1 Hz, 1H), 6.90 (d, *J* = 8.5 Hz, 1H), 4.91 (dd, *J* = 11.9, 5.3 Hz, 1H), 3.76 – 3.70 (m, 4H), 3.69 – 3.64 (m, 4H), 3.62 – 3.58 (m, 2H), 3.47 (t, *J* = 5.3 Hz, 2H), 2.90 – 2.65 (m, 3H), 2.15 – 2.07 (m, 1H).

**2-(2,6-dioxopiperidin-3-yl)-4-((2-(2-(2-(2-hydroxyethoxy)ethoxy)ethoxy)ethyl)amino)isoindoline-1,3-dione (8c):** Yellow oil, 30%. $^1$H NMR (400 MHz, DMSO-$d_6$) δ 11.08 (s, 1H), 7.63 – 7.55 (dd, *J* = 8.5, 7.0 Hz, 1H), 7.15 (d, *J* = 8.5 Hz, 1H), 7.04 (d, *J* = 7.0 Hz, 1H), 6.60 (t, *J* = 5.9 Hz, 1H), 5.05 (dd, *J* = 13.0, 5.4 Hz, 1H), 4.55 (t, *J* = 5.4 Hz, 1H), 3.62 (t, *J* = 5.3 Hz, 2H), 3.59 – 3.43 (m, 12H), 3.39 (t, *J* = 5.2 Hz, 2H), 2.94 – 2.82 (m, 1H), 2.56 (dd, *J* = 19.8, 10.4 Hz, 2H), 2.08 – 1.96 (m, 1H).

***Synthesis of compound P1-3*****:** compound **8a-c** (1.0 equiv) was dissolved in anhydrous DCM, DIPEA (2.0 equiv) and Cl-POCEN$^i$Pr$_2$ (1.5 equiv) was added. The mixture was stirred at room temperature for 1 hour. Solvent was removed, and the residue was purified with flash chromatography (Hexane:Actone (5%TEA)=100:0 to75:25), giving product **P1-3**.

**2-cyanoethyl (5-((2-(2,6-dioxopiperidin-3-yl)-1,3-dioxoisoindolin-4-yl)amino)pentyl) diisopropylphosphoramidite** (**P1**): Yellow oil, 65%. $^1$H NMR (400 MHz, DMSO-$d_6$) δ 11.08 (s, 1H), 7.57 (t, *J* = 7.9 Hz, 1H), 7.09 (d, *J* = 8.5 Hz, 1H), 7.01 (d, *J* = 6.2 Hz, 1H), 6.54 (s, 1H), 5.04 (dd, *J* = 12.4, 4.5 Hz, 1H), 3.78 –

3.65 (m, 2H), 3.64 – 3.45 (m, 4H), 2.95 – 2.82 (m, 1H), 2.74 (t, $J$ = 5.4 Hz, 2H), 2.63 – 2.52 (m, 2H), 2.02 (d, $J$ = 12.2 Hz, 1H), 1.59 (s, 4H), 1.42 (d, $J$ = 6.3 Hz, 2H), 1.15 (dt, $J$ = 13.9, 7.3 Hz, 12H).

**2-cyanoethyl (2-(2-(2-((2-(2,6-dioxopiperidin-3-yl)-1,3-dioxoisoindolin-4-yl)amino)ethoxy)ethoxy)ethyl) diisopropylphosphoramidite** (**P2**): Yellow oil, 68%. $^1$H NMR (400 MHz, DMSO-$d_6$) δ 11.08 (s, 1H), 7.61 – 7.54 (dd, $J$ = 8.6, 7.1 Hz, 1H), 7.14 (d, $J$ = 8.6 Hz, 1H), 7.04 (d, $J$ = 7.1 Hz, 1H), 6.60 (t, $J$ = 5.7 Hz, 1H), 5.05 (dd, $J$ = 12.9, 5.4 Hz, 1H), 3.79 – 3.66 (m, 2H), 3.61 (m, 2H), 3.59 – 3.50 (m, 10H), 3.47 (dd, $J$ = 11.0, 5.4 Hz, 2H), 2.88 (m, 1H), 2.75 (t, $J$ = 6.0 Hz, 2H), 2.63 – 2.52 (m, 2H), 2.06 – 1.99 (m, 1H), 1.12 (dd, $J$ = 6.7, 3.7 Hz, 12H).

**2-cyanoethyl (2-(2-(2-(2-((2-(2,6-dioxopiperidin-3-yl)-1,3-dioxoisoindolin-4-yl)amino)ethoxy)ethoxy)ethoxy)ethyl) diisopropylphosphoramidite (P3):** Yellow oil, 48%. $^1$H NMR (400 MHz, DMSO-$d_6$) δ 11.08 (s, 1H), 7.58 (dd, $J$ = 8.5, 7.2 Hz, 1H), 7.14 (d, $J$ = 8.6 Hz, 1H), 7.04 (d, $J$ = 7.0 Hz, 1H), 6.60 (t, $J$ = 5.7 Hz, 1H), 5.05 (dd, $J$ = 12.9, 5.4 Hz, 1H), 4.03 (m, 2H), 3.76 – 3.67 (m, 3H), 3.66 – 3.59 (m, 3H), 3.59 – 3.50 (m, 8H), 3.50 – 3.37 (m, 4H), 2.94 – 2.82 (m, 1H), 2.75 (t, $J$ = 6.0 Hz, 2H), 2.63 – 2.53 (m, 2H), 2.06 – 1.98 (m, 1H), 1.15 – 1.07 (m, 12H).

***Synthesis of compound 8d-f*:** Compound VHL-032 (1.0 equiv) was dissolved in DCM and DMF (1:1), and TEA (3.0 equiv), **7d-f** (1.5 equiv), and HATU (1.5 equiv) was added. The mixture was stirred at rt overnight. The reaction solution was diluted with DCM, washed with $NaHCO_3$ solution. The organic phase was concentrated and purified with flash chromatography (DCM:MeOH = 100:0 to 98:2), giving compound

**8d-f**.

**(2S,4R)-1-((S)-2-(6-((tert-butyldiphenylsilyl)oxy)hexanamido)-3,3-dimethylbutanoyl)-4-hydroxy-N-(4-(4-methylthiazol-5-yl)benzyl)pyrrolidine-2-carboxamide (8d) :** White foam solid, 70%. $^1$H NMR (400 MHz, $CDCl_3$) δ 8.72 (s, 1H), 7.64 (dd, *J* = 7.9, 1.6 Hz, 4H), 7.44 – 7.32 (m, 10H), 6.08 (d, *J* = 8.7 Hz, 1H), 4.70 (t, *J* = 7.9 Hz, 1H), 4.56 (dd, *J* = 15.0, 6.6 Hz, 1H), 4.49 (d, *J* = 8.8 Hz, 2H), 4.33 (dd, *J* = 15.0, 5.2 Hz, 1H), 4.11 – 4.05 (m, 1H), 3.61 (m, 3H), 2.57 – 2.49 (m, 4H), 2.16 (t, *J* = 7.6 Hz, 2H), 2.13 – 2.03 (m, 1H), 1.63 – 1.50 (m, 4H), 1.41 – 1.30 (m, 2H), 1.05 – 1.00 (m, 9H), 0.92 (s, 9H).

**(2S,4R)-1-((S)-14-(tert-butyl)-2,2-dimethyl-12-oxo-3,3-diphenyl-4,7,10-trioxa-13-aza-3-silapentadecan-15-oyl)-4-hydroxy-N-(4-(4-methylthiazol-5-yl)benzyl)pyrrolidine-2-carboxamide (8e):** Colorless oil, 62%. $^1$H NMR (400 MHz, $CDCl_3$) δ 8.72 (s, 1H), 7.69 – 7.63 (m, 4H), 7.44 – 7.28 (m, 10H), 4.73 (t, *J* = 7.9 Hz, 1H), 4.54 (m, 2H), 4.43 (d, *J* = 8.3 Hz, 1H), 4.32 (dd, *J* = 15.0, 5.3 Hz, 1H), 4.12 (d, *J* = 11.4 Hz, 1H), 3.99 (q, *J* = 15.8 Hz, 2H), 3.80 (dd, *J* = 7.8, 3.3 Hz, 2H), 3.71 – 3.54 (m, 7H), 2.56 (m, 1H), 2.51 (s, 3H), 2.14 – 2.06 (m, 1H), 1.06 – 1.00 (m, 9H), 0.92 (s,9H).

**(2S,4R)-1-((S)-17-(tert-butyl)-2,2-dimethyl-15-oxo-3,3-diphenyl-4,7,10,13-tetraoxa-16-aza-3-silaoctadecan-18-oyl)-4-hydroxy-N-(4-(4-methylthiazol-5-yl)benzyl)pyrrolidine-2-carboxamide (8f):** Colorless oil, 60%. $^1$H NMR (400 MHz, $CDCl_3$) δ 8.70 (s, 1H), 7.69 – 7.64 (m, 4H), 7.43 – 7.32 (m, 10H), 4.72 (t, *J* = 7.9 Hz, 1H), 4.53 (m, 2H), 4.47 (d, *J* = 8.5 Hz, 1H), 4.33 (dd, *J* = 15.0, 5.3 Hz, 1H), 4.08 (d, *J*

= 10.2 Hz, 1H), 4.03 – 3.91 (m, 2H), 3.79 (t, *J* = 5.3 Hz, 2H), 3.68 – 3.55 (m, 11H), 2.56 – 2.48 (m, 4H), 2.15 – 2.06 (m, 1H), 1.03 (d, *J* = 2.9 Hz, 9H), 0.94 (s, 9H).

***Synthesis of compound 9a-c*:** Compound **8d-f** (1.0 equiv) was dissolved in DCM and cooled to 0 ºC, then TEA (1.5 equiv) and DMAP (0.01 equiv) was added. The mixture was stirred and $Ac_2O$ (1.5 equiv) was added slowly. The reaction was stirred at 0ºC for 1h. the reaction solution was washed with water, and the organic phase was dried with $Na_2SO_4$, filtered and concentrated. The residue was purified with flash chromatography (DCM:MeOH = 100:0 to 98:2), giving compound **9a-c**.

**(3R,5S)-1-((S)-2-(6-((tert-butyldiphenylsilyl)oxy)hexanamido)-3,3-dimethylbutanoyl)-5-((4-(4-methylthiazol-5-yl)benzyl)carbamoyl)pyrrolidin-3-yl acetate (9a):** White foam solid, 90%.$^1$H NMR (400 MHz, $CDCl_3$) δ 8.89 (d, *J* = 3.7 Hz, 1H), 7.64 (dd, *J* = 7.6, 1.3 Hz, 4H), 7.43 – 7.32 (m, 10H), 7.18 – 7.13 (m, 1H), 6.04 (d, *J* = 9.1 Hz, 1H), 5.37 (s, 1H), 4.70 – 4.65 (m, 1H), 4.62 – 4.50 (m, 2H), 4.34 (dd, *J* = 14.9, 5.3 Hz, 1H), 4.05 (d, *J* = 12.7 Hz, 1H), 3.84 – 3.76 (m, 1H), 3.63 (t, *J* = 6.4 Hz, 2H), 2.71 (m, 1H), 2.54 (s, 3H), 2.17 (m, 3H), 2.03 (s, 3H), 1.57 (m, 4H), 1.36 (m, 2H),1.03 (s, 9H), 0.89 (s, 9H).

**(3R,5S)-1-((S)-14-(tert-butyl)-2,2-dimethyl-12-oxo-3,3-diphenyl-4,7,10-trioxa-13-aza-3-silapentadecan-15-oyl)-5-((4-(4-methylthiazol-5-yl)benzyl)carbamoyl)pyrrolidin-3-yl acetate (9b):** Colorless oil, 92%. $^1$H NMR (400 MHz, $CDCl_3$) δ 8.76 (s, 1H), 7.66 (dd, *J* = 7.8, 1.5 Hz, 4H), 7.43 – 7.30 (m, 10H), 7.22 (d, *J* = 8.4 Hz, 2H), 5.36 (s, 1H), 4.73 – 4.67 (m, 1H), 4.56 – 4.47 (m, 2H), 4.33 (dd, *J* = 14.9, 5.4 Hz, 1H), 4.05 (d, *J* = 11.9 Hz, 1H), 3.99 (d, *J* = 4.9 Hz, 2H),

3.84 – 3.75 (m, 3H), 3.70 – 3.56 (m, 7H), 2.77 – 2.69 (m, 1H), 2.52 (s, 3H), 2.15 (m, 1H), 2.03 (s, 3H), 1.03 (s, 9H), 0.90 (s, 9H).

**(3R,5S)-1-((S)-17-(tert-butyl)-2,2-dimethyl-15-oxo-3,3-diphenyl-4,7,10,13-tetraoxa-16-aza-3-silaoctadecan-18-oyl)-5-((4-(4-methylthiazol-5-yl)benzyl)carbamoyl)pyrrolidin-3-yl acetate (9c):** Colorless oil, 87%. $^1$H NMR (400 MHz, $CDCl_3$) δ 8.75 (s, 1H), 7.69 – 7.64 (m, 4H), 7.43 – 7.31 (m, 10H), 7.23 (dd, *J* = 14.1, 7.4 Hz, 2H), 5.36 (s, 1H), 4.71 (dd, *J* = 8.2, 6.6 Hz, 1H), 4.57 – 4.49 (m, 2H), 4.34 (dd, *J* = 14.9, 5.4 Hz, 1H), 4.05 (d, *J* = 13.7 Hz, 1H), 3.98 (d, *J* = 4.3 Hz, 2H), 3.80 (dd, *J* = 11.0, 5.8 Hz, 3H), 3.70 – 3.61 (m, 8H), 3.57 (t, *J* = 5.3 Hz, 2H), 2.77 – 2.68 (m, 1H), 2.52 (s, 3H), 2.20 – 2.13 (m, 1H), 2.04 (s, 3H), 1.06 – 1.01 (s, 9H), 0.91 (s, 9H).

***Synthesis of compound 10a-c*:** Compound **9a-c** (1.0 equiv) was dissolved in THF and TBAF (1M in THF, 2.0 equiv) was added. The mixture was stirred at rt overnight. The solvent was removed and the residue was purified with flash chromatography (DCM:MeOH = 100:0 to 97:3), giving compound **10a-c**.

**(3R,5S)-1-((S)-2-(6-hydroxyhexanamido)-3,3-dimethylbutanoyl)-5-((4-(4-methylthiazol-5-yl)benzyl)carbamoyl)pyrrolidin-3-yl acetate (10a):** White solid, 60%. $^1$H NMR (400 MHz, $CDCl_3$) δ 8.75 (s, 1H), 7.40 – 7.32 (m, 4H), 7.20 (t, *J* = 6.0 Hz, 1H), 6.03 (d, *J* = 9.2 Hz, 1H), 5.37 (m, 1H), 4.73 – 4.65 (m, 1H), 4.57 (dd, *J* = 14.9, 6.6 Hz, 1H), 4.51 (d, *J* = 9.2 Hz, 1H), 4.34 (dd, *J* = 14.9, 5.2 Hz, 1H), 4.07 (d, *J* = 11.7 Hz, 1H), 3.79 (dd, *J* = 11.6, 4.6 Hz, 1H), 3.66 – 3.57 (m, 2H), 2.75 – 2.66 (m, 1H), 2.54 (s, 3H), 2.19 (m, 3H), 2.05 (s, 3H), 1.64 (m, 2H), 1.60 – 1.51 (m, 2H), 1.47

(m, 2H), 0.90 (s, 9H).

**(3R,5S)-1-((S)-2-(2-(2-(2-hydroxyethoxy)ethoxy)acetamido)-3,3-dimethylbutanoyl)-5-((4-(4-methylthiazol-5-yl)benzyl)carbamoyl)pyrrolidin-3-yl acetate (10b):** White solid, 68%. $^{1}$H NMR (400 MHz, $CDCl_3$) δ 8.72 (s, 1H), 7.54 (d, $J$ = 9.5 Hz, 1H), 7.37 (s, 4H), 7.16 (t, $J$ = 5.8 Hz, 1H), 5.40 (m, 1H), 4.66 (dd, $J$ = 8.2, 6.7 Hz, 2H), 4.57 (dd, $J$ = 14.8, 6.6 Hz, 1H), 4.34 (dd, $J$ = 14.8, 5.4 Hz, 1H), 4.05 (dd, $J$ = 16.1, 5.5 Hz, 1H), 3.98 – 3.90 (m, 2H), 3.83 (dd, $J$ = 11.8, 4.7 Hz, 1H), 3.78 – 3.56 (m, 9H), 2.75 – 2.67 (m, 1H), 2.53 (d, $J$ = 3.3 Hz, 3H), 2.18 (m, 1H), 2.04 (d, $J$ = 2.5 Hz, 3H), 0.92 (s, 9H).

**(3R,5S)-1-((S)-2-(tert-butyl)-14-hydroxy-4-oxo-6,9,12-trioxa-3-azatetradecanoyl)-5-((4-(4-methylthiazol-5-yl)benzyl)carbamoyl)pyrrolidin-3-yl acetate (10c):** Colorless oil, 52%. $^{1}$H NMR (400 MHz, $CDCl_3$) δ 8.72 (s, 1H), 7.50 (dd, $J$ = 11.1, 5.3 Hz, 1H), 7.39 – 7.28 (m, 5H), 5.39 (m, 1H), 4.69 (dd, $J$ = 8.1, 6.4 Hz, 1H), 4.57 (m, 2H), 4.33 (dd, $J$ = 14.9, 5.3 Hz, 1H), 4.02 (d, $J$ = 8.6 Hz, 2H), 3.84 (dd, $J$ = 11.6, 4.9 Hz, 1H), 3.72 – 3.62 (m, 10H), 3.60 (m, 1H), 3.56 (m, 1H), 3.54 – 3.48 (m, 1H), 3.47 (d, $J$ = 1.4 Hz, 2H), 2.74 – 2.65 (m, 1H), 2.53 (d, $J$ = 4.0 Hz, 3H), 2.21 – 2.12 (m, 1H), 2.04 (s, 3H), 0.93 (s, 9H).

***Synthesis of compound P4-6*:** compound **10a-c** (1.0 equiv) was dissolved in anhydrous DCM, DIPEA (2.0 equiv) and Cl-POCEN$^{i}Pr_2$ (1.5 equiv) was added. The mixture was stirred at room temperature for 1 hour. Solvent was removed, and the residue was purified with flash chromatography (Hexane:Actone (5%TEA)=100:0 to 60:40), giving product as colorless oil.

**(3R,5S)-1-((2S)-2-(6-(((2-cyanoethoxy)(diisopropylamino)phosphaneyl)oxy)hexanamido)-3,3-dimethylbutanoyl)-5-((4-(4-methylthiazol-5-yl)benzyl)carbamoyl)pyrrolidin-3-yl acetate (P4):** Colorless oil, 60%. $^{1}$H NMR (400 MHz, $CDCl_3$) δ 8.68 (s, 1H), 7.36 (q, *J* = 8.1 Hz, 4H), 7.19 (t, *J* = 5.7 Hz, 1H), 6.01 (d, *J* = 9.1 Hz, 1H), 5.37 (m, 1H), 4.74 – 4.68 (m, 1H), 4.60 – 4.49 (m, 2H), 4.34 (dd, *J* = 14.7, 5.1 Hz, 1H), 4.04 (d, *J* = 12.1 Hz, 1H), 3.87 – 3.73 (m, 3H), 3.69 – 3.53 (m, 4H), 2.74 (m, 1H), 2.63 (t, *J* = 6.5 Hz, 2H), 2.52 (d, *J* = 0.6 Hz, 3H), 2.19 (m, 3H), 2.05 (s, 3H), 1.60 (m, 4H), 1.42 – 1.35 (m, 2H), 1.16 (q, *J* = 6.0 Hz, 12H), 0.89 (s, 9H).

**(3R,5S)-1-((2S)-2-(2-(2-(2-(((2-cyanoethoxy)(diisopropylamino)phosphaneyl)oxy)ethoxy)ethoxy)acetamido)-3,3-dimethylbutanoyl)-5-((4-(4-methylthiazol-5-yl)benzyl)carbamoyl)pyrrolidin-3-yl acetate (P5):** Colorless oil, 67%. $^{1}$H NMR (400 MHz, $CDCl_3$) δ 8.67 (s, 1H), 7.36 (q, *J* = 8.2 Hz, 4H), 7.26 – 7.22 (m, 1H), 7.19 (d, *J* = 9.2 Hz, 1H), 5.37 (m, 1H), 4.72 (dd, *J* = 8.0, 6.7 Hz, 1H), 4.59 – 4.48 (m, 2H), 4.35 (dd, *J* = 14.9, 5.3 Hz, 1H), 4.07 – 4.02 (m, 1H), 4.00 (d, *J* = 3.5 Hz, 2H), 3.91 – 3.76 (m, 4H), 3.75 – 3.64 (m, 7H), 3.59 ( m, 2H), 2.79 – 2.70 (m, 1H), 2.66 – 2.61 (m, 2H), 2.52 (s, 3H), 2.21 – 2.12 (m, 1H), 2.04 (s, 3H), 1.19 – 1.14 (m, 12H), 0.91 (s, 9H).

**(3R,5S)-1-((2S)-2-(tert-butyl)-14-(((2-cyanoethoxy)(diisopropylamino)phosphaneyl)oxy)-4-oxo-6,9,12-trioxa-3-azatetradecanoyl)-5-((4-(4-methylthiazol-5-yl)benzyl)carbamoyl)pyrrolidin-3-yl acetate (P6):** Colorless oil, 40%. $^{1}$H NMR (400 MHz, $CDCl_3$) δ 8.68 (s, 1H), 7.36 (q,

$J$ = 8.1 Hz, 4H), 7.25 – 7.17 (m, 2H), 5.37 (m, 1H), 4.75 – 4.69 (m, 1H), 4.59 – 4.49 (m, 2H), 4.36 (dd, $J$ = 14.9, 5.3 Hz, 1H), 4.07 – 4.02 (m, 1H), 4.00 (d, $J$ = 4.7 Hz, 2H), 3.90 – 3.75 (m, 4H), 3.75 – 3.53 (m, 13H), 2.80 – 2.71 (m, 1H), 2.64 (t, $J$ = 6.5 Hz, 2H), 2.52 (s, 3H), 2.16 (m, 1H), 2.04 (s, 3H), 1.21 – 1.14 (m, 12H), 0.92 (s, 9H).

**Synthesis of oligonucleotides**

All oligonucleotides used in this work were synthesized and reverse phase-HPLC purified by ExonanoRNA (Columbus, OH). Mass and purity (>95%) were confirmed by LC-MS from Novatia, LLC with Xcalibur system.

**Annealing reaction**

Single-stranded and reverse oligonucleotides were mixed in an assembly buffer (10 mM Tris-HCl [pH7.5], 100 mM NaCl, 1 mM EDTA), and heated to 90 $^{o}$C for 5 min, then slowly cool down to 37 $^{o}$C within 1 hour. Double-stranded O'PROTACs were mixed well, aliquoted and stored at -20 $^{o}$C for the future use.

**Cell culture and transfection**

VCaP, PC-3 and DU145 prostate cancer cell line and 293T cell line were obtained from the American Type Culture Collection (ATCC). 293T cells were maintained in DMEM medium with 10% FBS, PC-3 and DU145 cells were maintained in RPMI medium with 10% FBS. VCaP cells were cultured in RPMI medium with 15% FBS. Cells were transiently transfected using Lipofectamine 2000

(Thermo Fisher) for O’PROTAC according to the manufacturer’s instructions.

**Western blot**

Cell lysate was subjected to SDS-PAGE and proteins were transferred to nitrocellulose membranes (GE Healthcare Sciences). The membranes were blocked in Tris-buffered saline (TBS, pH 7.4) containing 5% non-fat milk and 0.1% Tween-20, washed twice in TBS containing 0.1% Tween-20, and incubated with primary antibody overnight at 4 °C, followed by secondary antibody for 1 hour at room temperature. The proteins of interest were visualized using ECL chemiluminescence system (Thermo Fisher).

**Biotin pull-down assay**

The 293T cells were transfected with 100 nM of biotin-labelled ERG O’ PROTACs and 1 μg of HA-ERG plasmid in 10-cm dishes using Lipofectamine 2000 (Thermo Fisher) for 36 h. The cells were treated with MG132 for 12 hours before lysed in lysis buffer containing 50 mM Tris-HCl (pH7.5), 150 mM NaCl, 1% NP-40, 0.5% sodium deoxycholate and 1% proteinase inhibitor. The cell lysate was incubated with Streptavidin Sepharose High Performance beads (GE Healthcare) overnight at 4 ºC. The binding protein was eluted by elution buffer and subjected to western blot.

**RNA extraction and RT-qPCR**

RNA was extracted using TRIzol (Invitrogen) and reversely transcribed into cDNA with SuperScript III First-Strand Synthesis System (Promega). The quantitative PCR (qPCR) was performed in the iQ thermal cycler (Bio-Rad) using the iQ SYBR Green Supermix (Bio-Rad). Each sample was carried out in triplicate and three biological repeats were performed. The ΔCT was calculated by normalizing the threshold difference of a certain gene with glyceraldehyde-3-phosphate dehydrogenase (*GAPDH*). The primer sequences are listed in Table S2.

**Cell growth assay**

PC-3 and DU145 cells were transfected with LEF1 OP-V1 for 48 hours and seeded in 96-well plate at the density of 1,000 per well. After cells adhered to the plate, at indicated time points, the CellTiter 96 Aqueous One solution Cell Proliferation Assay (MTS) (Promega) was added to each well to measure cell viability. MTS was diluted at a ratio of 1:10 in PBS and added into the wells and incubated for 2 hours at 37 °C in a cell incubator. Microplate reader was used to measure absorbance of 490 nm in each well.

**Table S1. The sequences of O’PROTACs**

| O’PROTAC | Sequence |
|---|---|
| ERG OP-C1 | Forward: 5'-ACGGACCGGAAATCCGGTT-3'<br>Reverse: 5'-P1-AACCGGATTTCCGGTCCGT-3' |
| ERG OP-C2 | Forward: 5'-ACGGACCGGAAATCCGGTT-3'<br>Reverse: 5'-P2-AACCGGATTTCCGGTCCGT-3' |

| | |
|---|---|
| ERG OP-C3 | Forward: 5'-ACGGACCGGAAATCCGGTT-3'<br>Reverse: 5'-P3-AACCGGATTTCCGGTCCGT-3' |
| ERG OP-V1 | Forward: 5'-ACGGACCGGAAATCCGGTT-3'<br>Reverse: 5'-P4-AACCGGATTTCCGGTCCGT-3' |
| ERG OP-V2 | Forward: 5'-ACGGACCGGAAATCCGGTT-3'<br>Reverse: 5'-P5-AACCGGATTTCCGGTCCGT-3' |
| ERG OP-V3 | Forward: 5'-ACGGACCGGAAATCCGGTT-3'<br>Reverse: 5'-P6-AACCGGATTTCCGGTCCGT-3' |
| FITC-ERG OP-C1 | Forward: 5'-FITC-ACGGACCGGAAATCCGGTT-3'<br>Reverse: 5'-P1-AACCGGATTTCCGGTCCGT-3' |
| Biotin-ERG OP-C1 | Forward: 5'-Biotin-ACGGACCGGAAATCCGGTT-3'<br>Reverse: 5'-P1-AACCGGATTTCCGGTCCGT-3' |
| LEF1 OP-C1 | Forward: 5'-TACAAAGATCAAAGGGTT-3'<br>Reverse: 5'-P1-AACCCTTTGATCTTTGTA-3' |
| LEF1 OP-C2 | Forward: 5'-TACAAAGATCAAAGGGTT-3'<br>Reverse: 5'-P2-AACCCTTTGATCTTTGTA-3' |
| LEF1 OP-C3 | Forward: 5'-TACAAAGATCAAAGGGTT-3'<br>Reverse: 5'-P3-AACCCTTTGATCTTTGTA-3' |
| LEF1 OP-V1 | Forward: 5'-TACAAAGATCAAAGGGTT-3'<br>Reverse: 5'-P4-AACCCTTTGATCTTTGTA-3' |
| LEF1 OP-V2 | Forward: 5'-TACAAAGATCAAAGGGTT-3'<br>Reverse: 5'-P5-AACCCTTTGATCTTTGTA-3' |

| | |
|---|---|
| LEF1 OP-V3 | Forward: 5'-TACAAAGATCAAAGGGTT-3'<br><br>Reverse: 5'-P6-AACCCTTTGATCTTTGTA-3' |

**Table S2. Primer sequence for RT-qPCR**

| Gene | Usage | Forward (5’- 3’) | Reverse (5’- 3’) |
|---|---|---|---|
| *ADAM19* | RT-qPCR | GCCTATGCCCCCTGAGAGTG | GCTTGAGTTGGCCTAGTTTGTTGTTC |
| *GAPDH* | RT-qPCR | GAAGGTGAAGGTCGGAGTC | GAAGATGGTGATGGGATTTC |
| *MMP3* | RT-qPCR | TTCATTTTGGCCATCTCTTCCTTCAG | TATCCAGCTCGTACCTCATTTCCTCT |
| *MMP9* | RT-qPCR | TGCCCGGACCAAGGATACAGT | AGCGCGTGGCCGAACTCAT |
| *PLAT* | RT-qPCR | CACTGGGCCTGGGCAAACATA | CACGTCAGCCTGCGGTTCTTC |
| *PLAU* | RT-qPCR | TACGGCTCTGAAGTCACCACCAAAAT | CCCCAGCTCACAATTCCAGTCAA |
